# Regulation of protein kinase pathways in primary neuron model of anti-NMDA receptor encephalitis

**DOI:** 10.64898/2026.09.15.751835

**Authors:** Yuyoung Joo, Prajwal Ciryam, Timothy Zhang, Weiliang Huang, Charles A. Dean, Audrey P. Lawrence, Thanh Hien Vu, Niki Gooya, Scott K. Dessain, Maureen A. Kane, David R. Benavides

## Abstract

The most common form of autoimmune encephalitis is associated with antibodies that target N-methyl-D-aspartic acid receptors (NMDARs). NMDARs play a pivotal role in neurotransmission and synaptic plasticity. Mounting evidence has shown that antibody targeting of the NMDAR GluN1 subunit, as in anti-NMDAR encephalitis, leads to NMDAR cross-linking and receptor internalization. However, the underlying signaling pathways affected by antibodies targeting NMDARs remain to be explored. We previously demonstrated that a human GluN1 monoclonal antibody 5F5 (GluN1 mAb 5F5) rapidly localizes to and regulates synaptic NMDAR function at native synapses of primary hippocampal neurons. Here, we sought to explore signaling targets of GluN1 mAb 5F5 in primary cortical neurons of either sex using subcellular fractionation, Western blotting, and label-free quantitative phosphoproteomics by mass spectrometry. We find that human GluN1 mAb 5F5 does not change NMDAR abundance or surface levels of GluN1 on primary cortical neurons at 2 hr. Despite this, we observe that GluN1 mAb 5F5 alters the phosphoproteome in synaptoneurosomes and regulates numerous synapse-related biological processes and protein kinase activities. Bioinformatic analyses suggest that these phosphoproteomic changes are positively correlated with NMDAR activation and negatively correlated with NMDAR inhibition. Together, these data suggest that GluN1 mAb 5F5 alters intracellular kinase signaling pathways in primary cortical neurons, likely by activating the NMDAR. These studies may help to identify novel therapeutic strategies for anti-NMDAR encephalitis and other antibody-mediated encephalitides targeting cell surface antigens.

**Significance statement:** Anti-NMDAR encephalitis is the most common form of autoimmune encephalitis. The pathophysiology of this clinical syndrome remains poorly defined but is currently thought to involve pathogenic antibodies that crosslink and internalize NMDARs. We identify intracellular signaling pathways targeted by human NMDAR antibodies in primary cortical neurons. Our key findings are obtained using specific, patient-derived human monoclonal antibodies with high affinity to the GluN1 subunit of NMDARs, which have been shown to induce key pathogenic effects of anti-NMDAR encephalitis (Sharma, Al-Saleem, Panzer, et al. 2018; Taraschenko, Fox, Eldridge, et al. 2021; Taraschenko, Fox, Zekeridou, et al. 2021; Dean et al. 2022). These studies expand the underlying pathophysiological molecular mechanisms involved in anti-NMDAR encephalitis.

## Introduction

Autoimmune encephalitis (AIE) is a severe brain disorder that results in poor recovery or death in up to 35% of patients (Broadley et al. 2019; Kvam et al. 2024). Cortical and hippocampal brain regions have been implicated in the pathogenesis of AIE (Kalaitzidis et al. 2023; Hartung, Cooper, et al. 2024; Hartung, von Schwanenflug, et al. 2024). Current treatments consist of nonspecific immunosuppression that does not directly address the consequences of antibodies on neuronal function. The lack of knowledge of the molecular neurobiology underlying AIE has slowed the development of disease-specific treatments.

Immunoglobulin G (IgG) antibody biomarkers are associated with AIE. The most common of these antibodies bind an extracellular epitope of the GluN1 subunit of N-methyl-D-aspartate receptors (NMDARs) causing anti-NMDAR encephalitis (Dalmau and Graus 2018). NMDARs are ionotropic glutamate receptors that govern excitatory synaptic transmission and NMDAR-dependent synaptic plasticity through control of calcium influx and postsynaptic signaling (Sabatini, Oertner, and Svoboda 2002; Lisman, Yasuda, and Raghavachari 2012). Cerebrospinal fluid IgGs to GluN1 are highly specific to anti-NMDAR encephalitis (Dalmau and Graus 2018; Hara et al. 2018) and bind to conserved extracellular epitopes in the amino terminal domain (GluN1-ATD) (Gleichman et al. 2012; Moscato et al. 2014; Michalski et al. 2024; Wang et al. 2024). The prevailing hypothesis is that GluN1 antibodies mediate their toxic effect by causing NMDAR crosslinking, internalization, and reduction of surface NMDARs at postsynaptic sites (Hughes et al. 2010; Moscato et al. 2014). GluN1 antibodies have been demonstrated, *in vitro* and *ex vivo*, to cause hippocampal NMDAR internalization, surface NMDAR reduction at postsynaptic sites, and diminished NMDAR-dependent long-term potentiation (LTP) (Mikasova et al. 2012; Planagumà et al. 2016; Planagumà et al. 2015). However, recent studies have described GluN1 antibodies that bind unique GluN1-ATD epitopes, regulate NMDAR channels, and fail to reduce NMDAR synaptic density (Wang et al. 2024; Michalski et al. 2024). One potentially powerful tool to understand the pathobiology of AIE are human patient-derived GluN1 monoclonal antibodies (GluN1 mAbs) (Kreye et al. 2016; Sharma, Al-Saleem, Panzer, et al. 2018; Malviya et al. 2017; Michalski et al. 2024; Wang et al. 2024). These antibodies have been interrogated in calcium responses and hippocampal physiology (Dean et al. 2022; Kreye et al. 2016; Malviya et al. 2017; Brahmer et al. 2010; Michalski et al. 2024). Previous work has utilized kinase microarray of lysates from primary hippocampal neurons exposed to human antibodies (Hunter et al. 2024) or proteomic analyses of hippocampal samples from a rodent passive transfer animal model (Ceanga et al. 2023) to explore molecular consequences of GluN1 antibodies. However, a unified model of the effects of GluN1 antibody exposure on intracellular signaling remains to be defined.

In the current study, we employed subcellular fractionation of primary cortical neurons to generate synaptoneurosomes (SNs) and postsynaptic density (PSD) fractions. SNs are enriched in presynaptic and postsynaptic elements relative to whole cell lysates. We isolated both SNs and PSD fractions after GluN1 mAb 5F5 exposure, for either 2 hr or 72 hr, to explore changes in synaptic signaling pathways. A quantitative phosphoproteomic analysis using mass spectrometry identified kinase signaling cascades regulated by GluN1 mAb 5F5 in SNs. We applied a suite of bioinformatic tools to characterize the effects of GluN1 mAb 5F5 on the phosphoproteome through pathway analysis, identification of putative upstream regulators, and comparison against published datasets in which NMDAR activity was modulated. Together, these analyses suggest that GluN1 mAb 5F5 acts through activation of the NMDAR. Further studies are warranted to dissect the exact mechanisms at play.

## Materials and Methods

### Animals

All experiments were performed with approval from the institutional Animal Care and Use Committee of University of Maryland School of Medicine.

### Rat primary neuron culture

Cortical tissue was dissected from embryonic day 18 (E18) Sprague-Dawley rat embryos, dissociated, and plated on poly-L-lysine-coated culture dishes in plating media (Neurobasal Plus medium supplemented with 2% B27 Plus, 2 mM GlutaMAX, 50 U/ml Penicillin-Streptomycin, and 5% FBS). Neurons were seeded at 65k/cm^2^ for biochemical analysis. At 4 days *in vitro* (DIV4), neurons were treated with 2 µM (+)-5-fluor-2’-deoxyuridine (FdU) for 24 hr, followed by exchange of growth media (Neurobasal Plus medium supplemented with 2% B27 Plus, 2 mM GlutaMAX, 50 U/ml Penicillin-Streptomycin). From DIV11 and every 3-4 days thereafter, cultures were supplemented with fresh growth media until DIV indicated. Neurons were used for biochemical analyses at DIV18-22.

### Human monoclonal antibodies

Human GluN1 mAbs were isolated from an 18 year-old female with anti-NMDAR encephalitis who presented with emotional lability, paranoia, and temporal lobe seizures (Sharma, Al-Saleem, Panzer, et al. 2018). GluN1 mAb 5F5 targets the GluN1 subunit of NMDAR in transfected HEK 293 cells (Sharma, Al-Saleem, Panzer, et al. 2018) as well as a cell line expressing GluN1-ATD (Sharma, Al-Saleem, Puligedda, et al. 2018). Control mAb 6A was used as IgG control for all experiments. Control mAb 6A is against BoNT serotype A (BoNT/A) heavy chain (Adekar et al. 2008) and is non-reactive in primary hippocampal neurons (Sharma, Al-Saleem, Panzer, et al. 2018; Dean et al. 2022). Primary cortical neurons treated with PBS served as vehicle control for all experiments. Human mAbs were incubated with neurons at a final concentration of 1 µg/ml in growth media for indicated times unless otherwise specified.

### Pharmacology

Primary cortical neurons were treated with vehicle (water) or NMDA (Tocris Bioscience, 0114) diluted in conditioned growth media at final concentration of 40 µM for 5 min at 37°C. Neurons were rinsed in ice-cold PBS and harvested for subcellular fractionation. In other experiments, primary cortical neurons were treated with vehicle (water) or (*RS*)-3,5-DHPG (Tocris Bioscience, 0342) diluted in conditioned growth media to final concentration of 100 µM for 5 min at 37°C followed by conditioned growth media exchange and 60 min incubation prior to harvest for surface biotinylation analyses.

### Subcellular fractionation of synaptoneurosomes and postsynaptic density

Primary cortical neurons were washed with ice-cold PBS and harvested in homogenization buffer 1 (HB1; 0.32 M sucrose and 4 mM HEPES, pH 7.4). The homogenates were disrupted mechanically using a 26G syringe, pushing the lysates up and down 10-12 times, and centrifuged at 800 x g for 10 min at 4°C. The soluble supernatant was collected from the nuclear pellet and centrifuged at 17,000 x g for 20 min at 4°C. The supernatant was collected as cytosol fraction. For SN fraction, the second pellet was washed in HB1, centrifuged at 17,000 x g for 20 min at 4°C, and resuspended in appropriate buffer as SN fraction. For PSD fraction preparation, the second pellet was lysed by hypoosmotic shock by resuspending in ice-cold deionized water (DW).

HEPES was immediately added to the final concentration of 4 mM, pH 7.4. The lysate was incubated at 4°C with rotation for 30 min and centrifuged by ultracentrifuge at 24,600 x g for 20 min at 4°C. After resuspending the pellets in a homogenization buffer 2 (HB2: 2 mM EDTA, 50 mM HEPES, pH 7.4), Triton X-100 solution was added to final concentration of 0.5%. After incubation for 15 min at 4°C with rotation, the lysates were ultracentrifuged at 32,557 x g for 20 min at 4°C. The final pellet (PSD fraction) was collected and resuspended in RIPA buffer, consisting of (in mM): Tris-HCl (pH 7.5) 20, NaCl 150, Na_2_EDTA 1, EGTA 1, NP-40 1%, sodium deoxycholate 1%, sodium pyrophosphate 2.5, β-glycerophosphate 1, Na_3_VO_4_ 1, and 1 µg/ml leupeptin. HB1, HB2, DW and RIPA buffer were supplemented with protease and phosphatase inhibitors, including 1 mM Na_3_VO_4_, Phosphatase Inhibitor Cocktail I (Sigma), Phosphatase Inhibitor Cocktail III (Sigma), and Roche cOmplete, EDTA-free protease inhibitor cocktail. Protein concentrations were quantified using bicinchoninic acid (BCA) protein assay (Thermo Scientific).

### Western blotting and quantification

Protein concentrations of all samples were determined using BCA protein assay and equal amounts of total protein (10 µg) from each sample were resolved by SDS-PAGE and transferred to PVDF membranes. REVERT total protein stain was utilized on all membranes as loading control (LI-COR Biosciences). Western blotting was carried out using standard protocols (Metzbower et al. 2019) utilizing near-infrared fluorescence labeled secondary antibodies. Western blot images were obtained on Odyssey CLx Infrared Imaging System and quantitated using Image Studio 4.0 software (LI-COR Biosciences). Protein abundance values were calculated relative to total protein stain (REVERT, LI-COR Biosciences) for each lane in all cases prior to comparative analyses. Protein phosphoprotein levels were calculated by dividing phosphorylated signal by the target protein signal prior to comparative analyses. Quantitative plots were constructed using normalized mean values from samples collected from each condition across experiments. Data represent mean values +/-standard error of the mean (SEM). Statistical analyses were conducted on quantitated values using GraphPad Prism software.

### Surface biotinylation assay

Surface biotinylation was performed as previously reported (Diering, Gustina, and Huganir 2014) with slight modifications. Neurons were rinsed with ice-cold PBS, pH 8.0, containing 0.1 mM CaCl_2_ and 1 mM MgCl_2_ (PBSCM) and incubated with Sulfo-NHS-SS-biotin (Thermo Scientific) at 0.5 mg/ml in PBSCM for 30 min at 4°C. Neurons were rinsed in PBSCM and unreacted biotinylation reagent was quenched twice in PBSCM containing 20 mM glycine for 7 min at 4°C. Neurons were lysed in lysis buffer (PBS containing 50 mM NaF, 5 mM sodium pyrophosphate, 1 mM EDTA, 1% IGEPAL CA-630, 0.5% sodium deoxycholate, 0.02% SDS, 1 µM okadaic acid, 1 mM Na_3_VO_4_, and Roche cOmplete, EDTA-free protease inhibitor cocktail) for 20 min at 4°C, scraped and transferred to microcentrifuge tubes. Protease and phosphatase inhibitors were added to buffers immediately before use. Lysates were centrifuged at 17,000 x g for 15 min at 4°C and supernatants were collected. The protein concentration of each lysate was quantified using BCA protein assay, and equal amounts of protein were incubated with NeutrAvidin coupled agarose beads (Thermo Scientific) overnight at 4°C. The beads were washed with ice-cold lysis buffer, and biotinylated proteins were eluted with 2x Laemmli sample buffer (BioRad). Total or surface samples were then subjected to SDS-PAGE and analyzed by Western blot.

### Mass spectrometry and differential abundance estimation

SN fractions were prepared from primary cortical neurons (DIV20-21) as described above (omitting protease inhibitors) and resuspended in 0.32 M sucrose and 4 mM HEPES/DW, pH 7.4. Differential expression of proteins were determined using high-resolution liquid chromatography-tandem mass spectrometry (LC-MS/MS) using label-free quantitation like our previous studies (Shah et al. 2023; Zalesak-Kravec et al. 2022; Martinez et al. 2022; Nelson et al. 2021; Huang et al. 2021; Huang, Yu, Liu, Tudor, et al. 2020; Huang, Yu, Liu, Defnet, et al. 2020; Chan et al. 2020; Defnet et al. 2019; Kim et al. 2019; Nelson et al. 2019; Huang, Yu, Jones, Carter, Jackson, et al. 2019; Huang, Yu, Jones, Carter, Pierzchalski, et al. 2019). The samples were enriched in phosphopeptides (relative to whole cell lysate) by affinity chromatography during which most nonphosphopeptides were depleted. SN samples were washed in PBS and solubilized in 4% sodium deoxycholate, reduced, alkylated, and digested with trypsin after which tryptic peptides were analyzed using a Waters nanoACQUITY UPLC coupled to a ThermoScientific Orbitrap Fusion Lumos Tribrid mass spectrometer. Tandem mass spectra were searched against a UniProt Mus musculus reference proteome using a Sequest HT algorithm and a MS Amanda algorithm described previously (Eng et al. 2008; Dorfer et al. 2014) with max precursor mass error tolerance of 10 ppm. Identifications were validated at a maximum FDR of 0.01 using a semi-supervised machine learning algorithm Percolator (The et al. 2016; Granholm et al. 2014). Label-free quantifications were performed using Minora, an aligned AMRT cluster quantification algorithm (Horn et al. 2016). Protein abundance ratios between groups were measured by comparing the MS1 peak volumes of peptide ions, whose identities were confirmed by MS2 sequencing. Imputation was performed to replace missing phosphopeptide values with reasonable estimates based on the available data, limiting calculated abundance ratios to max of 1000 or min of 0.001. All p-values reported were adjusted for multi-testing by applying a Benjamini-Hochberg correction procedure (Benjamini and Hochberg 1995). Phosphopeptides with *Q* value < 0.05 and abs(log_2_ fold change) ≥ 0.585 were considered differentially expressed. UpSetR was used to visualize comparisons between multiple datasets (Conway, Lex, and Gehlenborg 2017).

### Principal component analysis, pathway analysis, and Gene Ontology (GO)

Prior to performing principal component analysis (PCA), we imputed missing values using the function MinProb with q = 0.01 in the R package DEP (Zhang et al. 2018). PCA and Spearman’s correlation on the phosphopeptides were performed in R. We searched using DAVID (Database for Annotation, Visualization, and Integrated Discovery) bioinformatics tool and did not find any overrepresented pathways. QIAGEN Ingenuity Pathway Analysis (IPA, version 01-23-01) was used to identify putative overrepresented pathways and upstream regulators inferred from differential phosphorylation at the primary phosphorylated site (Site 1) between GluN1 mAb 5F5 and Control mAb 6A at 2 hr and 72 hr after treatment. Thresholds were as noted above (adjusted p < 0.05, abs(log_2_ fold change) ≥ 0.585) and default settings for IPA Core Analysis were used. Upstream Analysis was performed to identify regulators as activated or inhibited based on abs(activation Z-score) ≥ 2 and overlap p < 0.05. Those flagged by IPA as biased results were excluded. We performed semi-automated overrepresentation analysis for Gene Ontology (GO) terms using the *elim* method in the R package topGO (Alexa, Rahnenführer, and Lengauer 2006).

### Kinase motif overrepresentation analysis

Only singly phosphorylated phosphopeptides were included in the analysis. Abundance values for identical peptide sequences were combined based on identity irrespective of non-phosphorylation post-translational modifications. Differential expression analysis was conducted in R as previously reported using the Differential Enrichment analysis of Proteomics data package (Zhang et al. 2018). Phosphopeptides missing observations in all replicates were excluded, while the remaining missing values were imputed using random values from a Gaussian distribution centered around a minimal value, as previously reported (Huber et al. 2002). The differential expression-based analysis for kinase motif overrepresentation (Johnson et al. 2023) was conducted using PhosphoSitePlus (https://phosphosite.org), using the Fisher Enrichment Analysis differential phosphorylation method. The overrepresentation analysis method used was Percentile Rank, with default settings (enrichment threshold = 15, adjusted p < 0.1). The threshold for differentially expressed peptides used was adjusted p < 0.05 and abs(log_2_ fold change) ≥ 0.585.

### Comparative analysis to JB2 NMDAR activator

An IGFBP2-mimetic peptide fragment, JB2, functions as an NMDAR activator (Burgdorf et al. 2023). Raw data from rat primary cortical neurons treated at DIV21 for 1 hr with vehicle or 1 mM JB2 were filtered for Ser/Thr residue modifications (Burgdorf et al. 2023). Odds ratios and Holm-Bonferroni-adjusted p-values were calculated to quantify the overlap between proteins differentially phosphorylated in response to GluN1 mAb 5F5 and those differentially phosphorylated in response to JB2. For kinase motif overrepresentation analysis using PhosphoSitePlus, settings were the same as above, except that the threshold for differentially expressed peptides used was abs(log_2_ fold change) ≥ 1, as set by the authors of that study (Burgdorf et al. 2023).

### Experimental design and statistical analyses

Quantitated phosphoprotein levels were normalized against total protein levels. Quantitative plots were constructed using average values from 3-10 experiments. Statistical analysis was conducted on quantitated values using GraphPad software (GraphPad Prism 10.0, Boston, Massachusetts USA). All quantitative data are depicted as mean ± SEM. Comparisons between two groups were made using two-tailed Student’s t-test. For comparisons between more than two groups, a one-way analysis of variance (ANOVA) was used with appropriate post-hoc test. Two-way ANOVA was used to examine the effects of human mAb and drug on protein levels, with Tukey’s multiple comparison test performed as post-hoc test. Two-way ANOVA was used to examine the effects of human mAb and time on protein levels, with Tukey’s multiple comparison test performed as post-hoc test. All biochemical experiments were repeated at least three times. In all analyses, a *p* < 0.05 was considered statistically significant, unless otherwise stated.

### Antibodies

Primary antibodies used in this study include rabbit polyclonal GluN1 C-terminus (1:2,000, Sigma-Aldrich Cat# G8913, RRID:AB_259978); rabbit polyclonal GluN2A N-terminus (1:500, JH6097 gift from R. Huganir); mouse monoclonal GluN2B C-terminus (1:2,000, Millipore Cat# 05-920, RRID:AB_417391); mouse monoclonal PSD-95 (1:5,000, UC Davis/NIH NeuroMab Facility Cat# 75-028, RRID:AB_2292909); rabbit monoclonal synaptophysin (1:40,000, Abcam Cat# ab32127, RRID:AB_2286949); mouse monoclonal β-tubulin (Sigma-Aldrich Cat# T8578, RRID:AB_1841228); mouse monoclonal calcium/calmodulin-dependent protein kinase II (CaMKII) α (1:2,000, Thermo Fisher Scientific Cat# MA1-048, RRID:AB_325403); mouse monoclonal CaMKIIβ (1:5,000, Thermo Fisher Scientific Cat# 13-9800, RRID:AB_2533045); rabbit polyclonal pThr286 CaMKII (1:5,000, Cell Signaling Technology Cat# 3361, RRID:AB_10015209); mouse monoclonal Akt (1:1,000, Cell Signaling Technology Cat# 58295, RRID:AB_2799545); mouse monoclonal pThr308 Akt (1:1,000, Cell Signaling Technology Cat# 5106, RRID:AB_836861); rabbit monoclonal pSer473 Akt (1:1,000, Cell Signaling Technology Cat# 4060, RRID:AB_2315049). Near-infrared conjugated secondary antibodies (LI-COR Biosciences, Lincoln, Nebraska USA) were used to detect Western blot signals.

### CODE ACCESSIBILITY

Analysis code available upon request.

## Results

### Effect of anti-NMDAR antibodies on NMDAR subunit and PSD-95 levels in primary neurons

The binding characteristics of GluN1 mAb 5F5, isolated from a patient with anti-NMDAR encephalitis, on primary hippocampal neurons have been reported previously (Sharma, Al-Saleem, Panzer, et al. 2018; Sharma, Al-Saleem, Puligedda, et al. 2018; Dean et al. 2022). We have previously shown GluN1 mAb 5F5 demonstrates synaptic labeling in primary neurons, consistent with synaptic NMDAR targeting (Dean et al. 2022; Sharma, Al-Saleem, Panzer, et al. 2018). Having established that GluN1 mAb 5F5 targets primary neurons, we sought to evaluate the synaptic content of NMDARs and PSD-95 at the biochemical level following human mAb exposure and NMDAR activation. To accomplish this, we incubated primary cortical neurons with vehicle, Control mAb 6A, or GluN1 mAb 5F5 for 2 hr. A schematic of the experimental design is shown (Fig. 1A). To activate NMDARs, we used pharmacological activation *in vitro* with NMDA (40 µM) for 5 min prior to subcellular fractionation. We then interrogated NMDAR subunit levels (i.e., GluN1, GluN2A, and GluN2B) and PSD-95 levels via quantitative Western blotting. We observed that synaptoneurosomes (SNs) were enriched in presynaptic (i.e., synaptophysin) and postsynaptic (i.e., GluN1 and PSD-95) proteins compared to crude homogenates (Fig. 1B). Further, we observed increases in PSD proteins, and reduction of synaptophysin, in PSD fractions compared to crude homogenates (Fig. 1B). Therefore, we focused our studies on SN and PSD fractions (Fig. 1C).

**Figure 1:**
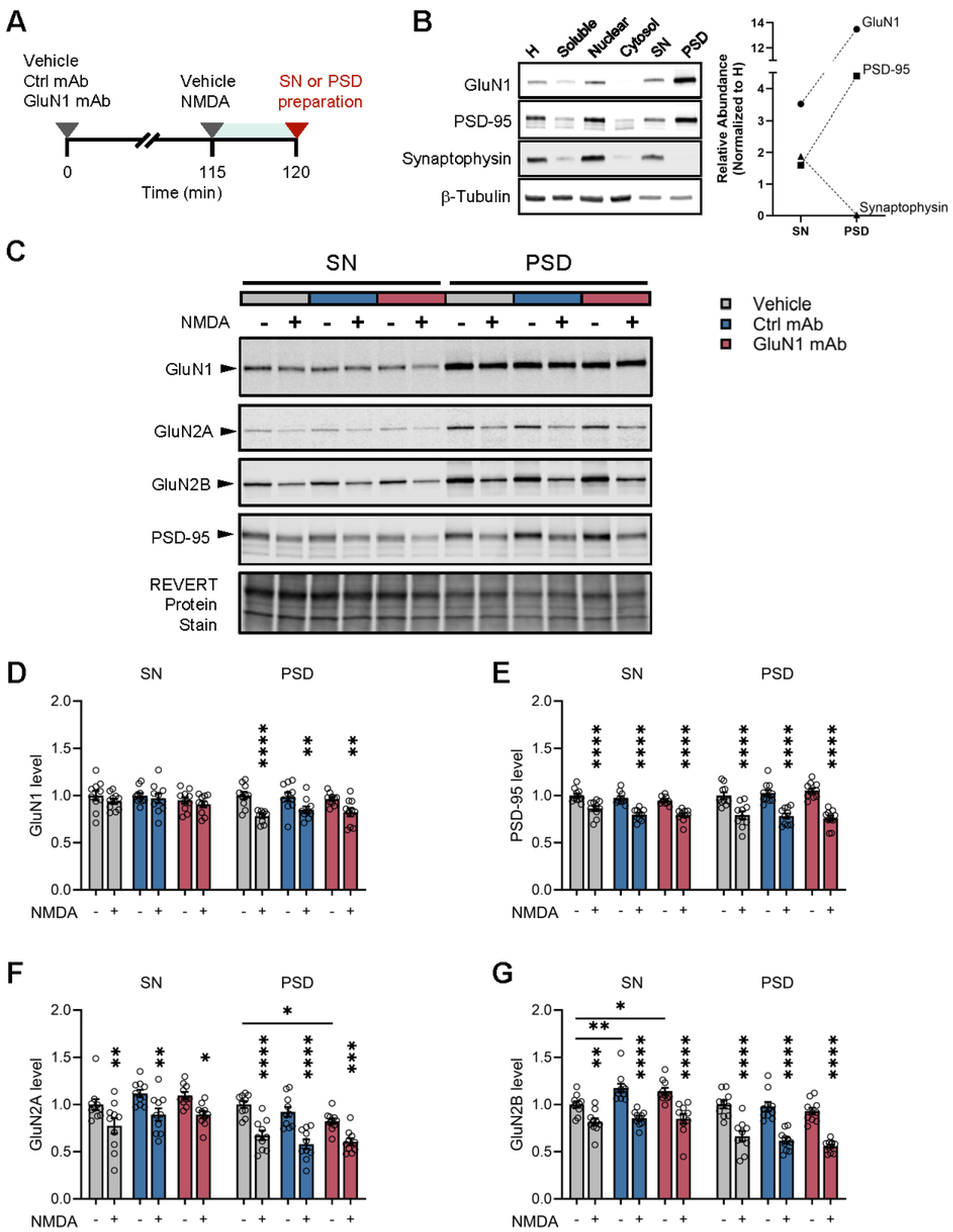
Human GluN1 mAb does not alter NMDAR levels in primary cortical neurons. A, Schematic experimental timeline for human mAb exposure and pharmacology. B, Western blot validation of subcellular fractions from primary cortical neurons. Representative blots for GluN1, PSD-95, synaptophysin, and β-tubulin are shown. Quantification of normalized relative abundance is shown for GluN1 (circle), PSD-95 (square), and synaptophysin (triangle). All targets are enriched in SN compared to H, while only GluN1 and PSD-95 are enriched in PSD. C, Representative Western blots for GluN1, GluN2A, GluN2B, and PSD-95 are shown from primary cortical neurons following vehicle (grey), Control mAb (blue), and GluN1 mAb (red) exposure. Representative image of REVERT total protein stain is also shown. D, Quantification of GluN1 levels in SN and PSD of primary cortical neurons following vehicle, Control mAb, and GluN1 mAb exposure. No effect of human mAb or NMDA on GluN1 levels was detected in SN fractions. NMDA reduces GluN1 levels in PSD in vehicle (0.78 versus 1.00 no NMDA, *p* < 0.0001), Control mAb (0.85 versus 0.99 no NMDA, *p* = 0.0098) and GluN1 mAb (0.82 versus 0.96 no NMDA, *p* = 0.0077) conditions. n = 10 replicates. Data are mean ± SEM, with individual replicates. E, Quantification of PSD-95 levels following Ab and NMDA. NMDA reduces PSD-95 levels in SN in vehicle (0.86 versus 1.00 no NMDA, *p* < 0.0001), Control mAb (0.80 versus 0.97 no NMDA, *p* < 0.0001) and GluN1 mAb (0.79 versus 0.95 no NMDA, *p* < 0.0001) conditions. NMDA reduces PSD-95 levels in PSD in vehicle (0.79 versus 1.00 no NMDA, *p* < 0.0001), Control mAb (0.78 versus 1.03 no NMDA, *p* < 0.0001) and GluN1 mAb (0.76 versus 1.05 no NMDA, *p* < 0.0001) conditions. n = 10 replicates. Data are mean ± SEM, with individual replicates. F, Quantification of GluN2A levels in SN and PSD. NMDA reduces GluN2A levels in SN in vehicle (0.77 versus 1.00 no NMDA, *p* = 0.0058), Control mAb (0.89 versus 1.12 no NMDA, *p* = 0.0064) and GluN1 mAb (0.89 versus 1.10 no NMDA, *p* = 0.0122) conditions. GluN1 mAb reduces GluN2A levels compared to vehicle in PSD (*p* = 0.0167). NMDA reduces GluN2A levels in PSD in vehicle (0.67 versus 1.00 no NMDA, *p* < 0.0001), Control mAb (0.58 versus 0.93 no NMDA, *p* < 0.0001) and GluN1 mAb (0.61 versus 0.82 no NMDA, *p* = 0.0008) conditions. n = 10 replicates. Data are mean ± SEM, with individual replicates. G, Quantification of GluN2B levels in SN and PSD. Control mAb and GluN1 mAb increase GluN2B levels compared to vehicle in SN (*p* = 0.0082 and *p* = 0.0364, respectively). NMDA reduces GluN2B levels in SN in vehicle (0.82 versus 1.00 no NMDA, *p* = 0.0017), Control mAb (0.86 versus 1.17 no NMDA, *p* < 0.0001) and GluN1 mAb (0.85 versus 1.14 no NMDA, *p* < 0.0001) conditions. NMDA reduces GluN2B levels in PSD in vehicle (0.67 versus 1.00 no NMDA, *p* < 0.0001), Control mAb (0.62 versus 0.98 no NMDA, *p* < 0.0001) and GluN1 mAb (0.56 versus 0.93 no NMDA, *p* < 0.0001) conditions. n = 10 replicates. Data are mean ± SEM, with individual replicates. * *p* < 0.05 (two-way ANOVA plus Tukey’s test), ** *p* < 0.01 (two-way ANOVA plus Tukey’s test), *** *p* < 0.001 (two-way ANOVA plus Tukey’s test), **** *p* < 0.0001 (two-way ANOVA plus Tukey’s test). H, homogenate; SN, synaptoneurosome; PSD, postsynaptic density.

Given that some GluN1 antibodies act in part by promoting receptor internalization, we first sought to establish the effect of GluN1 mAb 5F5 on the density of NMDAR subunits and associated proteins in the synapse. As a positive control, we used NMDA stimulation, which is well-established to alter surface receptor density through activity-dependent receptor trafficking and internalization mechanisms (Roche et al. 2001; Ehlers 2003). As expected, NMDA reduced the levels of GluN1 (*p* < 0.0001), PSD-95 (*p* <0.0001), GluN2A (*p* < 0.0001), and GluN2B (*p* < 0.0001) in the PSD (Fig. 1D-G). NMDA also reduced the levels of PSD-95 (*p* < 0.0001), GluN2A (*p* = 0.0058), and GluN2B (*p* = 0.0017), but not GluN1 (*p* = 0.168p value) in the SN. Surprisingly, we did not see similar effects for GluN1 mAb 5F5. Compared to Control mAb 6A, GluN1 mAb 5F5 did not have an effect on GluN1, PSD-95, GluN2A, or GluN2B in either SN or PSD (Fig. 1D-G). GluN1 mAb 5F5 reduced GluN2A levels in PSD samples compared to vehicle (*p* = 0.017), but no difference compared to Control mAb 6A. Simple main effect analysis showed that antibody had a statistically significant effect on SN GluN2B levels. Post-hoc comparisons showed increased SN GluN2B levels in both Control mAb (*p* = 0.0082) and GluN1 mAb (*p* = 0.0364) treatment compared to vehicle (Fig. 1G). However, no difference was detected between GluN1 mAb 5F5 compared to Control mAb. We next tested whether GluN1 mAb 5F5 might act to modify the effect of NMDA on synaptic protein levels. However, two-way ANOVA did not reveal statistically significant interactions between antibody and NMDA on GluN1, PSD-95, GluN2A, or GluN2B levels in SN or PSD samples. Together, these data suggest subtle differences in GluN1 mAb 5F5 regulation of GluN2A content in PSD samples when compared to vehicle condition. These findings may be consistent with a prior report that nanostructures increase at 2 hr following antibody exposure (Ladépêche et al. 2018). However, the GluN1 mAb 5F5 changes reported here were not statistically different from Control mAb 6A.

We wondered whether GluN1 mAb 5F5 might alter surface NMDAR levels, despite only a modest effect on total levels in SN and PSD fractions. At 2 hr, GluN1 mAb 5F5 did not alter surface GluN1 levels compared to controls as determined by surface biotinylation assay (Figure S1A). No effect on total GluN1 levels, nor surface GluN1 levels, were observed in any group compared to untreated samples. As a positive control (Figure S1B), the surface biotinylation assay was validated using primary cortical neurons treated with DHPG (100 µM, 5 min), which led to reduced GluN1 surface level (0.80 versus 1.00 vehicle, *p* = 0.0157), as expected (Palmer et al. 1997; Snyder et al. 2001). Together, these findings suggest that synaptic NMDAR levels and surface expression are unaffected by GluN1 mAb 5F5 in primary cortical neurons at 2 hr. These findings suggest that the early effects of GluN1 mAb 5F5 in primary cortical neurons may extend beyond regulation of surface NMDAR levels and instead involve other mechanisms governing NMDAR signaling and function.

### Phosphoproteomics and differentially expressed phosphoproteins following anti-NMDAR antibodies

NMDARs modulate synaptic plasticity through control of calcium influx and postsynaptic signaling by a robust intracellular network of pathways (Sabatini, Oertner, and Svoboda 2002; Lisman, Yasuda, and Raghavachari 2012). Having observed the effect of 2 hr incubation with GluN1 mAb 5F5 on NMDAR subunit levels in SN fractions, we assessed the state of intracellular signaling cascades using quantitative mass-spectrometry based phosphoproteomic analysis. If changes were observed at 2 hr, this would support the interpretation that GluN1 mAb 5F5 alters downstream functions independent of changes in NMDAR surface expression. To determine the effect of GluN1 mAb 5F5 on protein phosphorylation signaling cascades, we treated primary cortical neurons with GluN1 mAb 5F5 or controls and prepared SN fractions for label-free quantitative mass-spectrometry and bioinformatics analysis (Fig. 2A). Treatment conditions consisted of vehicle, Control mAb 6A, or GluN1 mAb 5F5 incubated for 2 hr or 72 hr prior to isolation of SN fractions. Using this approach, we detected 8379 peptides derived from 2271 unique proteins. Most of the observed peptides were phosphorylated (6268/8379, 75%), indicating the overall efficiency of phosphopeptide enrichment (relative to cell lysate) and detection (Data S1). Our method of affinity chromatography would be expected to enrich for peptides that are phosphorylated on Ser/Thr residues. Consistent with this, a phosphorylation motif analysis indicated that the most common phosphorylation sites identified in our dataset were primarily proline-directed Ser/Thr residues (Figure S2A).

**Figure 2:**
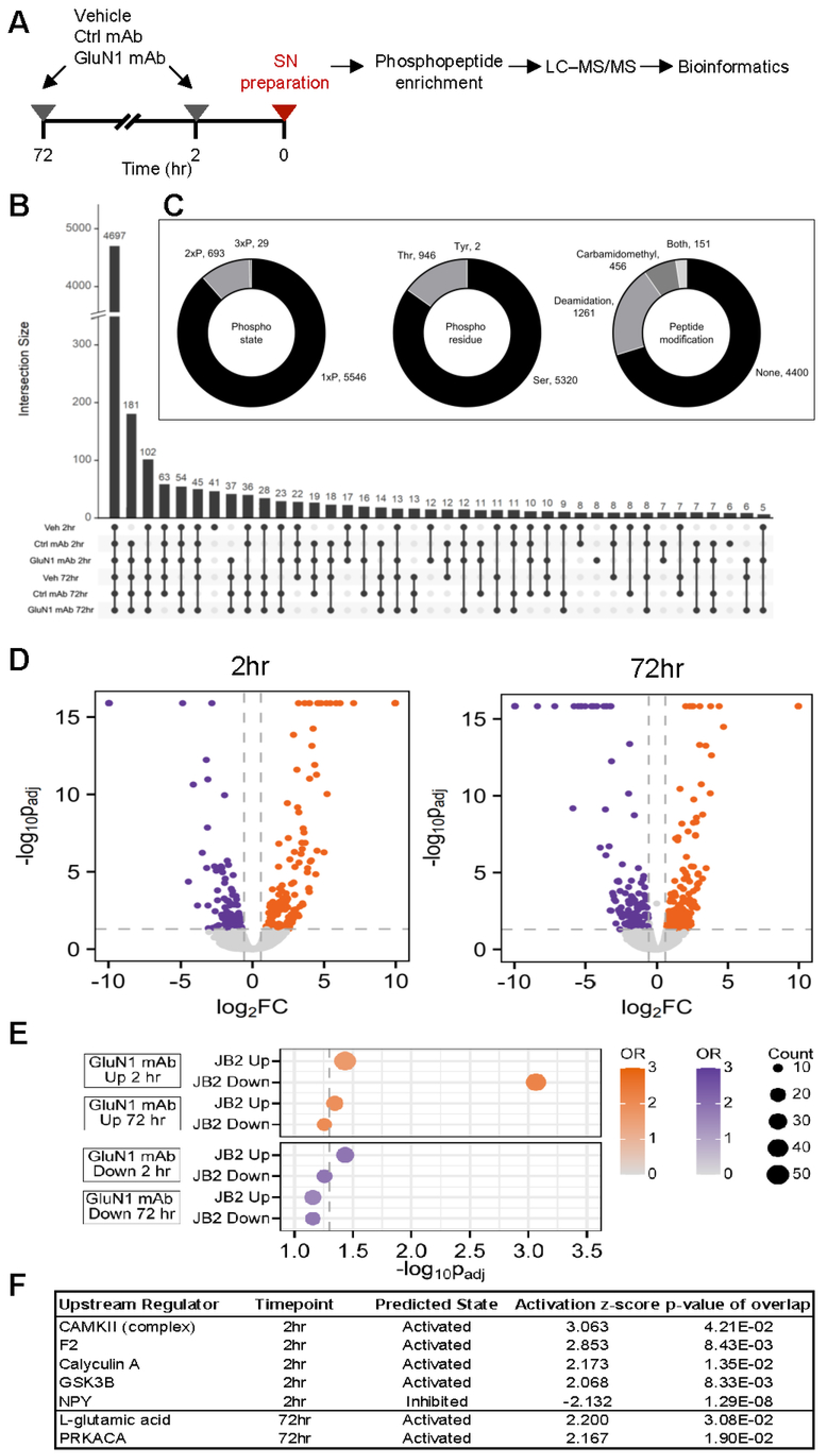
Modulation of protein phosphorylation by human GluN1 mAb in synaptoneurosomes. A, Summary of the experimental procedure. Rat primary cortical neurons were treated with Veh, Ctrl mAb, or GluN1 mAb on DIV19 or DIV21 and were harvested at the same time. SN were prepared and submitted for quantitative, label-free phosphoproteomics and bioinformatic analysis. B, Upset plot of phosphopeptide distribution across different treatment conditions and times. Most phosphopeptides (n = 4697) were found in all groups. The Veh 2 hr condition had the most unique phosphopeptides (n = 41). C, Phosphorylation state, phosphorylation residue, and protein modifications present on identified phosphopeptides. Most detected phosphopeptides are singly phosphorylated (5546/6268), with Ser being the most common modified residue (5320/6268). A minority of phosphopeptides also contain either deamidated, carbamidomethylated, or both modification on some residues (1868/6268). D, Volcano plot of differentially expressed phosphopeptides in GluN1 mAb condition at 2 hr and 72 hr timepoints. Only Ser/Thr kinases were included (n = 311). Cutoffs used for phosphosite differential expression were *p* < 0.05 and log_2_FC ≥ 0.585. E, Dotplot comparing differentially expressed phosphoproteins for GluN1 mAb (versus Ctrl mAb) to differentially expressed phosphoproteins for NMDAR activator JB2 (versus vehicle). Those proteins whose phosphopeptides were upregulated by JB2 are denoted “JB2 Up” while those who phosphopeptides were downregulated by JB2 are denoted “JB2 Down.” F, Results of IPA Upstream Analysis to identify putative upstream regulators inferred from differential phosphorylation at the primary phosphorylated site (Site 1) between GluN1 mAb 5F5 and Control mAb 6A at 2 hr and 72 hr after treatment. Thresholds were adjusted p < 0.05, abs(log_2_ fold change) ≥ 0.585 and default settings for IPA Core Analysis were used. Upstream Analysis was performed to identify regulators as activated or inhibited based on abs(activation Z-score) ≥ 2 and overlap p < 0.05. Those flagged by IPA as biased results were excluded.

Most phosphopeptides (4697/6268, 75%) were detected in all conditions (Fig. 2B), indicating consistent coverage across experimental conditions. Further, most phosphopeptides contained a single phosphorylation, which occurred most often on a Ser residue (Fig. 2C). Some deamidated and carbamidomethylated peptides were observed, but most phosphopeptides (4400/6268, 70%) were otherwise unmodified (Fig. 2C). Spearman’s rank correlation coefficient showed a positive linear correlation in phosphopeptides between treatment conditions (Figure S2B), the most closely related biological replicates showing p = 0.93 (Figure S2C). Overall, these findings are consistent with high data quality of representative phosphopeptides from SN fractions of primary neurons.

PCA revealed that first and second PCs accounted for 21.9% and 18.2% of the variance of the data respectively (Figure S2D). As an indication of data quality, biological replicates mapped to similar regions on the biplot and we observed separation between groups based on condition. We noted that variance between Control mAb 6A and GluN1 mAb 5F5 was most evident in PC2, while PC1 largely reflected variation of all other samples, particularly Veh 2 hr. It is unclear why the Veh 2 hr condition was highly dissimilar to the other groups (Figure S2D) and contained the most unique phosphopeptides amongst all conditions (Fig. 2B). We reasoned that including the Veh conditions as the control in our analyses would therefore skew the results. Phosphopeptides that were uniquely regulated by GluN1 mAb would be of particular interest for further characterization. Therefore, to avoid confounding comparisons and conclusions, we compared only the Control mAb and GluN1 mAb conditions in a revised subset analysis. Here, two PCs were responsible for 48.2% variance (Figure S2E), with excellent separation between groups based on condition.

We sought to define differentially regulated phosphopeptides across treatment conditions. As some peptides are not observed in all conditions (Fig. 2B), we imputed maximum ratios for comparisons in which one condition had a missing value. Differentially expressed phosphopeptides between GluN1 mAb 5F5 and Control mAb 6A condition were determined by ANOVA (FDR < 0.05) (Fig. 2D). At 2 hr, we found increased abundance of 384 phosphopeptides derived from 313 proteins and decreased abundance of 219 phosphopeptides derived from 179 proteins. At 72 hr, we found increased abundance of 160 phosphopeptides derived from 143 proteins and decreased abundance of 208 phosphopeptides derived from 172 proteins.

### Upstream regulation of the phosphoproteome is consistent with GluN1 mAb-induced activation of the NMDAR

We hypothesized that the GluN1 mAb mediates its effect on the phosphoproteome, at least in part, by agonizing the NMDAR. The resemblance of the GluN1 mAb-induced phosphoproteome to that arising from NMDAR activation motivated us to compare differentially regulated lists to a previously published study evaluating an NMDAR activator in SNs from primary cortical neurons (Burgdorf et al. 2023). IGFBP2 has been shown to enhance excitatory synaptic transmission and promote LTP by enhancing NMDAR function (Khan et al. 2019). An IGFBP2-derived peptide, JB2, induced changes in protein phosphorylation in synaptic membrane phosphoproteome at 1 hr (Burgdorf et al. 2023). We reasoned that if the GluN1 mAb 5F5 modulated NMDAR activation, then there should be significant overlap between the JB2-induced phosphoproteome and the GluN1 mAb-induced phosphoproteome. To test this, we mapped the differentially expressed phosphopeptides at each timepoint following GluN1 mAb to their precursor proteins and compared these lists against the precursor proteins of the phosphopeptides differentially expressed in response to JB2. We identified significant overlap in GluN1 mAb 5F5 regulated proteins compared to JB2 treatment (Fig. 2E), particularly at the 2 hr timepoint and not the 72 hr timepoint. Given the acute effects of JB2 on NMDARs, this result suggests that changes induced by GluN1 mAb at 2 hr are consistent with altered NMDAR function, while changes at 72 hr may reflect homeostatic or compensatory changes following NMDAR internalization. We note that proteins with phosphosites upregulated by GluN1 mAb 5F5 at 2 hr significantly overlap with those upregulated and downregulated by JB2 (Fig. 2E). Taken together, these data suggest that GluN1 mAb 5F5 induces extensive remodeling of the phosphoproteome consistent with altered NMDAR function.

One limitation of this approach is that we looked at the protein level instead of the phosphopeptide level, in part to avoid overinterpreting the JB2-induction data set, which included a single sample per condition. We reasoned that the effect of GluN1 mAb 5F5 on NMDAR function might best be understood by considering phosphopeptide changes. For this reason, we undertook an unbiased analysis to identify regulators of the phosphoproteome through which the GluN1 mAb 5F5 might act. By elucidating regulators associated with the GluN1 mAb-induced phosphoproteome signature, we sought to clarify whether GluN1 mAb behaved in a fashion that resembled activation at the NMDAR. We used the Upstream Analysis function in the IPA platform. We identified a set of regulators with a predicted activation state of *activated* or *inhibited* (Fig. 2F). Activated regulators are those whose expected effect on the phosphoproteome is positively correlated with the changes we observed in the phosphoproteome, while the effect of inhibited regulators is negatively correlated with our data. We found four activated regulators at 2 hr (CaMKII, F2, calyculin A, GSK3β); one inhibited regulator at 2 hr (Neuropeptide Y); and 2 activated regulators at 72 hr (l-glutamic acid, PRKACA) (Data S2). The strongest effect based on Z-score (3.063) was observed for CaMKII at 2 hr. CaMKII is highly abundant in the brain and is a key downstream mediator of NMDAR function in LTP. GSK3β is a kinase that helps to stabilize NMDAR at the membrane (Amici et al. 2021). F2 (prothrombin) is a precursor of thrombin, which activates the protease-activated receptor-1 (PAR1), a potentiator of NMDAR (Gingrich et al. 2000). Calyculin A is a phosphatase inhibitor that has been shown to enhance NMDAR activation (Blank et al. 1997). Neuropeptide Y, an inhibitor of NMDAR (Fu, Wessel, and Taylor 2020), is negatively correlated with our phosphoproteome signature at 2 hr. Conversely, l-glutamic acid, an agonist of NMDARs, is positively correlated with our phosphoproteome signature at 72 hr. PRKACA is the catalytic subunit of Protein Kinase A, which plays a critical role in NMDAR activation (Skeberdis et al. 2006). These regulators act through a wide variety of mechanisms both upstream and downstream of NMDARs. The correlation (both positive and negative) of each of these regulators with the GluN1 mAb-induced phosphoproteome suggests that GluN1 mAb 5F5 acts as an activator of the NMDAR.

### Gene Ontology reveals synaptic, cytoskeletal, and signaling processes overrepresented in the GluN1 mAb-induced phosphoproteome

We next sought to identify biological processes (BP) and molecular functions (MF) regulated by GluN1 mAb compared to Control mAb. We mapped the differentially regulated phosphopeptides at each timepoint to their precursor protein and gene to conduct GO term overrepresentation analysis. As previously indicated, we focused on phosphopeptides that were uniquely regulated by GluN1 mAb 5F5 compared to Control mAb 6A at 2 hr timepoint, to emphasize GO BP and MF governed by GluN1 mAb 5F5. Functional overrepresentation analysis identified many significantly overrepresented GO BP and MF terms (Data S3). Of these, 18 GO BP and 16 GO MF terms were amongst the top 5 most significantly overrepresented in at least one condition (Fig. 3).

**Figure 3:**
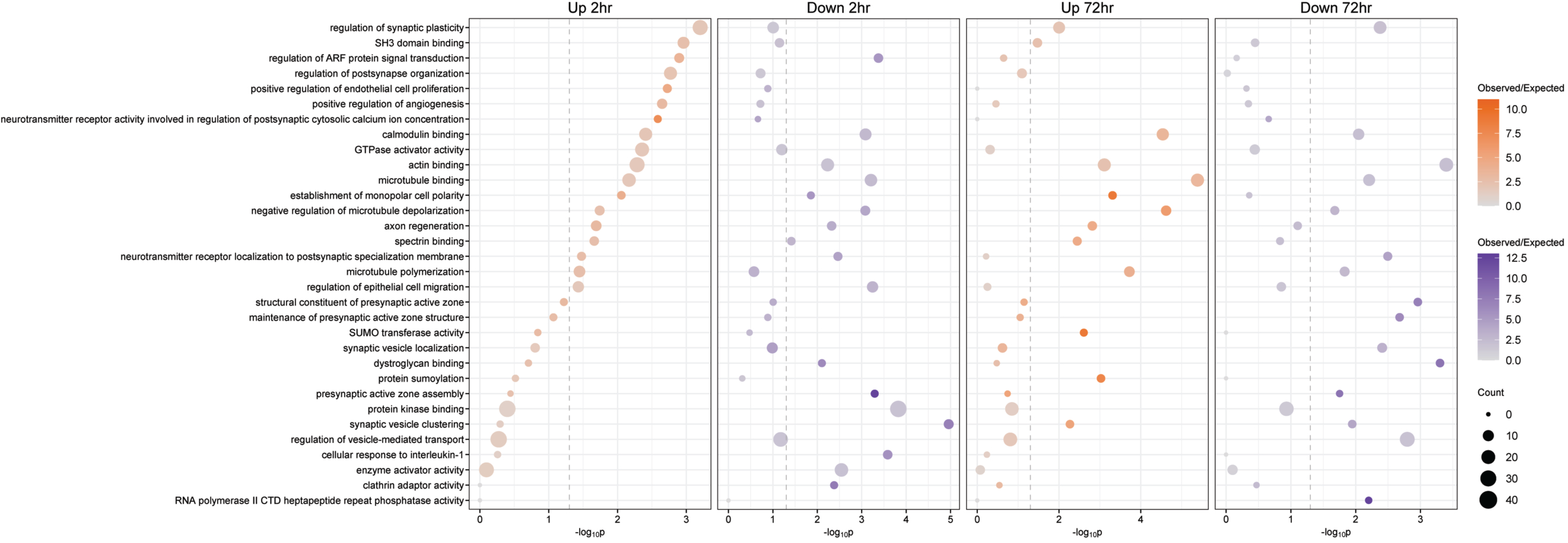
Functional overrepresentation analysis of Gene Ontology biological processes and molecular functions by human GluN1 mAb. Dotplot of selected biological processes and molecular function GO terms identified by topGO as being overrepresented among genes that correspond to differentially expressed phosphosites in the GluN1 mAb 5F5 condition. The plotted terms are those that were in the top five most overrepresented among either upregulated or downregulated phosphopeptides with GluN1 treatment at either 2h or 72h. The full list of all genes in dataset was used as the background for overrepresentation analysis.

Among the overrepresented terms in the upregulated lists at 2 hr were GO BP “regulation of synaptic plasticity” and “regulation of postsynapse organization” and GO MF term “neurotransmitter receptor activity involved in regulation of postsynaptic cytosolic calcium ion concentration.” Critical signaling GO MF terms of “SH3 domain binding,” “GTPase activator activity,” and “actin binding” were upregulated at 2 hr, suggesting an effect on actin remodeling in the early timepoints following GluN1 mAb 5F5 exposure. Interestingly, GO MF “calmodulin binding” and “regulation of ARF protein signal transduction” were top terms in both the upregulated and downregulated lists at 2 hr exposure. These may suggest nuanced regulation of these signaling pathways. Interestingly, downregulated lists at 2 hr were associated with “synaptic vesicle clustering” and “presynaptic active zone assembly”, implying regulation of presynaptic function. Overall, these data support the interpretation that GluN1 mAb 5F5 exposure alters cellular behaviors critical to the structure and function of neuronal synapses.

We also evaluated the most overrepresented GO BP and MF terms following 72 hr GluN1 mAb exposure (Fig. 3). The most overrepresented GO BP terms in upregulated list at 72 hr include “negative regulation of microtubule depolymerization,” “microtubule polymerization” and GO MF term “microtubule binding.” Furthermore, the most overrepresented GO BP terms in the downregulated list at 72 hr include “regulation of vesicle-mediated transport,” and “neurotransmitter receptor localization to postsynaptic specialization membrane.” These findings are consistent with the prevailing observation that GluN1 antibodies cause NMDAR internalization and reduction at postsynaptic sites (Hughes et al. 2010; Moscato et al. 2014). Interestingly, “protein sumoylation” and “SUMO transferase activity” are top terms in upregulated lists at 72 hr, suggesting these processes may be involved in GluN1 mAb-induced NMDAR trafficking (Berndt, Wilkinson, and Henley 2012). Furthermore, top terms in downregulated lists at 72 hr include “regulation of vesicle-mediated transport,” “neurotransmitter receptor localization to postsynaptic specialization membrane,” and “regulation of synaptic plasticity.” Together, these data support the interpretation that GluN1 mAb 5F5 alters protein phosphorylation cascades, biological processes, and molecular functions critical to the structure and function of neuronal synapses that change over time.

### Evaluation of calmodulin-regulated pathways by anti-NMDAR antibodies

Calmodulin-related pathways featured prominently in our bioinformatic analysis. CaMKII was the top hit from IPA Upstream Analysis. “Calmodulin binding” was also one of only four of the top GO pathways overrepresented in proteins with both upregulated and downregulated phosphopeptides at 2 hr and 72 hr (the others were related to the cytoskeleton). This prompted us to investigate CaMKII regulation by GluN1 mAb 5F5 at 2 hr. Our phosphoproteomic data showed that phosphorylation state of CaMKII was differentially regulated at 2 hr and 72 hr timepoints, but many CaMKII phosphopeptides either had multiple modifications and/or could not be mapped due to technical limitations. Similar peptides were detected separately that may represent an MS artifact. To avoid related bias, we used the data from all detected phosphopeptides. In the phosphoproteomic dataset, phosphorylation sites that were regulated by GluN1 mAb 5F5 included Ser330, Ser330/Ser331, Ser331/T334 CaMKIIα, and Ser394/Thr401 CaMKIIβ (Data S1). These CaMKII residues map to the variable linker region of the molecule (Fig. 4A), whose phosphorylation is not as well characterized as that in the regulatory region. Reliable phospho-specific antibodies to detect these sites are unavailable. Therefore, we undertook validation experiments by testing canonical CaMKII phosphorylation sites that were not significantly changed by GluN1 mAb. A schematic of the experimental design is shown (Fig. 4B).

**Figure 4:**
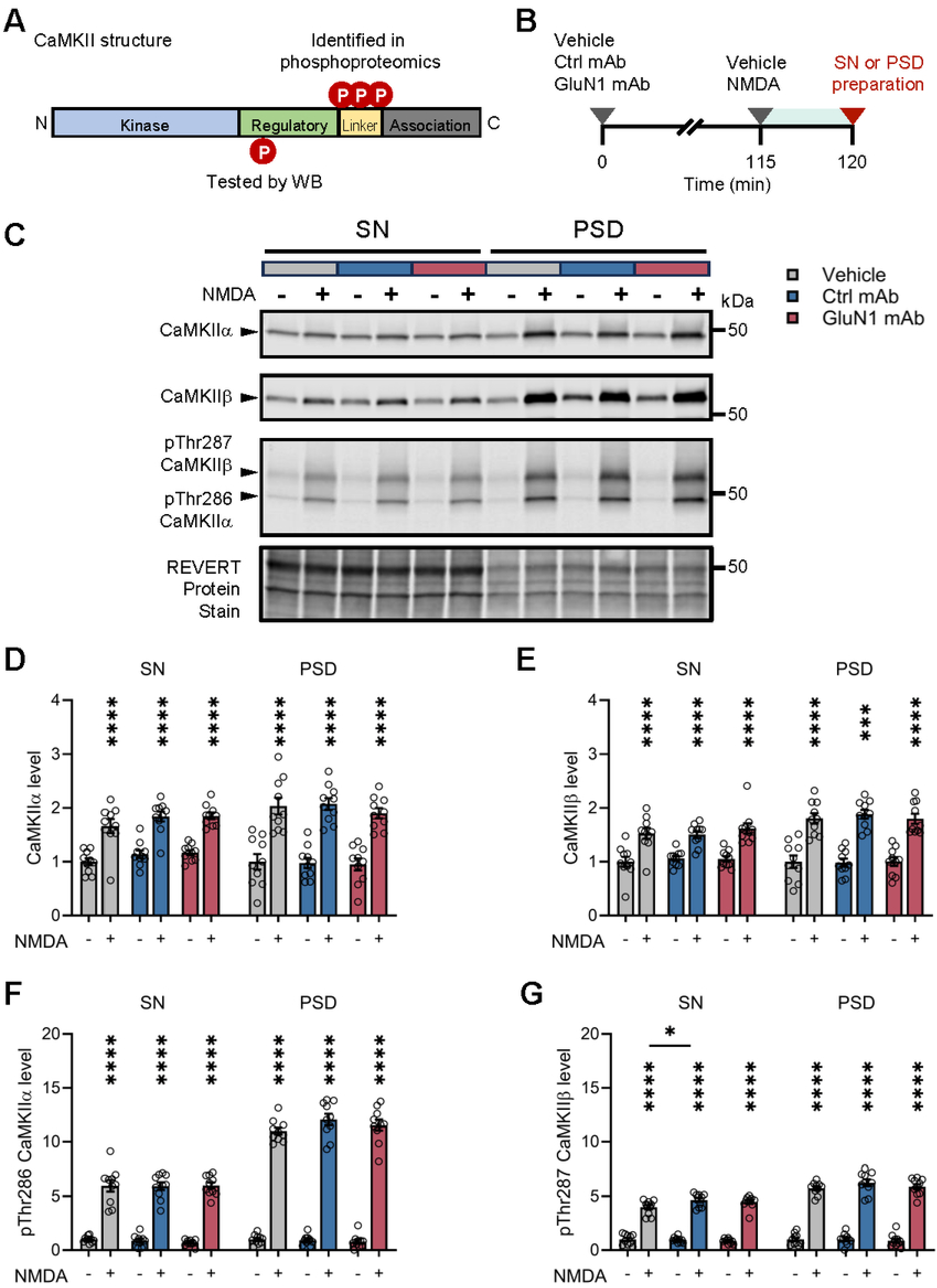
Evaluation of CaMKII in primary cortical neurons following human GluN1 mAb exposure. A, Schematic of CaMKII isozyme structure demonstrating kinase, regulatory, linker, and association domains. Location of phosphosites identified in phosphoproteomics dataset are shown. Phosphorylation of Thr286 CaMKIIα (Thr287 CaMKIIβ) in the regulatory domain, which we tested by Western blotting (WB), is responsible for keeping the kinase autonomous of calcium/calmodulin and partially active (∼20%–40% maximal activity) (Miller and Kennedy 1986; Coultrap et al. 2010). B, Schematic experimental timeline for human mAb exposure and pharmacology. C, Representative Western blots for CaMKIIα, CaMKIIβ, and pThr286 CaMKIIα (pThr287 CaMKIIβ) are shown from primary cortical neurons following vehicle (grey), Control mAb (blue), and GluN1 mAb (red) exposure. Representative image of REVERT total protein stain is also shown. D, Quantification of CaMKIIα levels in SN and PSD of primary cortical neurons following vehicle, Control mAb, and GluN1 mAb exposure. No effect of human mAb on CaMKIIα levels was detected in SN or PSD fractions. NMDA increases CaMKIIα levels in SN in vehicle (1.66 versus 1.00 no NMDA, *p* < 0.0001), Control mAb (1.84 versus 1.14 no NMDA, *p* < 0.0001) and GluN1 mAb (1.85 versus 1.17 no NMDA, *p* < 0.0001) conditions. NMDA increases CaMKIIα levels in PSD in vehicle (2.03 versus 1.00 no NMDA, *p* < 0.0001), Control mAb (2.07 versus 0.98 no NMDA, *p* < 0.0001) and GluN1 mAb (1.90 versus 0.95 no NMDA, *p* < 0.0001) conditions. n = 10 replicates. Data are mean ± SEM, with individual replicates. E, Quantification of CaMKIIβ levels in SN and PSD of primary cortical neurons following vehicle, Control mAb, and GluN1 mAb exposure. There was no effect of human mAb on CaMKIIβ levels in SN or PSD fractions. NMDA increases CaMKIIβ levels in SN in vehicle (1.53 versus 1.00 no NMDA, *p* < 0.0001), Control mAb (1.50 versus 1.06 no NMDA, *p* = 0.0002) and GluN1 mAb (1.62 versus 1.05 no NMDA, *p* < 0.0001) conditions. NMDA increases CaMKIIβ levels in PSD in vehicle (1.81 versus 1.00 no NMDA, *p* < 0.0001), Control mAb (1.89 versus 0.98 no NMDA, *p* < 0.0001) and GluN1 mAb (1.80 versus 1.00 no NMDA, *p* < 0.0001) conditions. n = 10 replicates. Data are mean ± SEM, with individual replicates. F, Quantification of pThr286 CaMKIIα levels in SN and PSD of primary cortical neurons following vehicle, Control mAb, and GluN1 mAb exposure. No effect of human mAb on pThr 286 CaMKIIα levels was detected in SN or PSD fractions. NMDA increases pThr286 CaMKIIα levels in SN in vehicle (5.98 versus 1.00 no NMDA, *p* < 0.0001), Control mAb (5.94 versus 0.90 no NMDA, *p* < 0.0001) and GluN1 mAb (6.00 versus 0.70 no NMDA, *p* < 0.0001) conditions. NMDA increases pThr286 CaMKIIα levels in PSD in vehicle (10.98 versus 1.00 no NMDA, *p* < 0.0001), Control mAb (12.06 versus 0.93 no NMDA, *p* < 0.0001) and GluN1 mAb (11.55 versus 0.79 no NMDA, *p* < 0.0001) conditions. n = 10 replicates. Data are mean ± SEM, with individual replicates. G, Quantification of pThr287 CaMKIIβ levels in SN and PSD. There was no effect of human mAb on CaMKIIβ levels. NMDA increases pThr287 CaMKIIβ levels in SN in vehicle (3.99 versus 1.00 no NMDA, *p* < 0.0001), Control mAb (4.65 versus 0.96 no NMDA, *p* < 0.0001) and GluN1 mAb (4.48 versus 0.85 no NMDA, *p* < 0.0001) conditions. There was a significant effect of NMDA in pThr287 CaMKIIβ levels in SN fractions between Control mAb and vehicle conditions (*p* = 0.0118). NMDA increases pThr287 CaMKIIβ levels in PSD in vehicle (5.75 versus 1.00 no NMDA, *p* < 0.0001), Control mAb (6.26 versus 1.01 no NMDA, *p* < 0.0001) and GluN1 mAb (5.90 versus 0.88 no NMDA, *p* < 0.0001) conditions. n = 10 replicates. Data are mean ± SEM, with individual replicates. * *p* < 0.05 (two-way ANOVA plus Tukey’s test), ** *p* < 0.01 (two-way ANOVA plus Tukey’s test), *** *p* < 0.001 (two-way ANOVA plus Tukey’s test), **** *p* < 0.0001 (two-way ANOVA plus Tukey’s test). H, homogenate; SN, synaptoneurosome; PSD, postsynaptic density.

Activation of calcium/calmodulin signaling results in CaMKII autophosphorylation at Thr286 (pThr286) CaMKIIα, corresponding to pThr287 CaMKIIβ, which generates an autonomously active form of the kinase (Miller and Kennedy 1986; Lou, Lloyd, and Schulman 1986; Schworer, Colbran, and Soderling 1986), independent of calcium/calmodulin (Fig. 4B). To explore this, we utilized subcellular fractionation of primary neurons and quantitative Western blotting to evaluate CaMKII isoform (i.e., CaMKIIα and CaMKIIβ) and autophosphorylation levels (i.e., pThr286 CaMKIIα or pThr287 CaMKIIβ) following antibody treatment and NMDA stimulation (Fig. 4C). Acute NMDAR activation by NMDA (40 µM, 5 min) resulted in an increase in CaMKII localization in the SN and PSD fractions of both CaMKII isoforms (Fig. 4D,E), as well as robust increase in levels of autophosphorylation of CaMKIIα and CaMKIIβ in all conditions (Fig. 4F,G). These control experiments established our ability to detect phosphorylation changes at these residues in our system. By contrast, GluN1 mAb 5F5 did not demonstrate a significant effect on phosphorylation levels of pThr286 CaMKIIα and pThr287 CaMKIIβ (Fig. 4D-G). There was not a statistically significant interaction between the effects of antibody or NMDA on CaMKII levels or autophosphorylated CaMKII levels in SN or PSD fractions (Fig. 4). These specific observations for CaMKII are consistent with those derived from phosphoproteomics. These data suggest that GluN1 mAb 5F5 dysregulates CaMKII signaling in a manner distinct from NMDA.

Calmodulin activity is also a critical upstream regulator of Akt (Protein Kinase B). Akt expression changes have been shown in mutated calmodulin proteins (Cheng et al. 2003; Zheng et al. 2008), is critical in synaptic plasticity and microtubule dynamics (Itoh et al. 2016; Read and Gorman 2009), and a major negative regulator of GSK3 (Cross et al. 1995; Dudek et al. 1997). Therefore, we sought to explore the regulation of Akt following GluN1 mAb 5F5 exposure (Fig. 5). To do so, we used time course experiments in primary neurons to determine effects of GluN1 mAb 5F5 and Control mAb 6A compared to vehicle conditions in cytosolic fractions. This analysis revealed total Akt levels increased following human mAb exposure (Fig. 5B). However, there was no difference between Control mAb and GluN1 mAb conditions in post hoc analyses. Next, we utilized quantitative Western blotting to evaluate activating phosphorylation levels on Akt (i.e., pThr308 or pSer473) following antibody treatment (Fig. 5C,D). These phosphosites were not present in our phosphoproteomic dataset (Data S1). Simple main effects analyses showed that time had a statistically significant effect on both pThr308 (F(2,60) = 2.78, p = 0.0489) and pSer473 (F(2,60) = 33.88, p < 0.0001) levels. Additionally, a simple main effect of antibody exposure had a statistically significant effect on pSer473 (F(2,60) = 3.88, p = 0.0260) (Fig. 5D). However, there were no differences between Control mAb and GluN1 mAb conditions in post hoc analyses on pThr308 or pSer473 levels. These data suggest that the Akt activation is not altered following exposure to GluN1 mAb 5F5.

**Figure 5:**
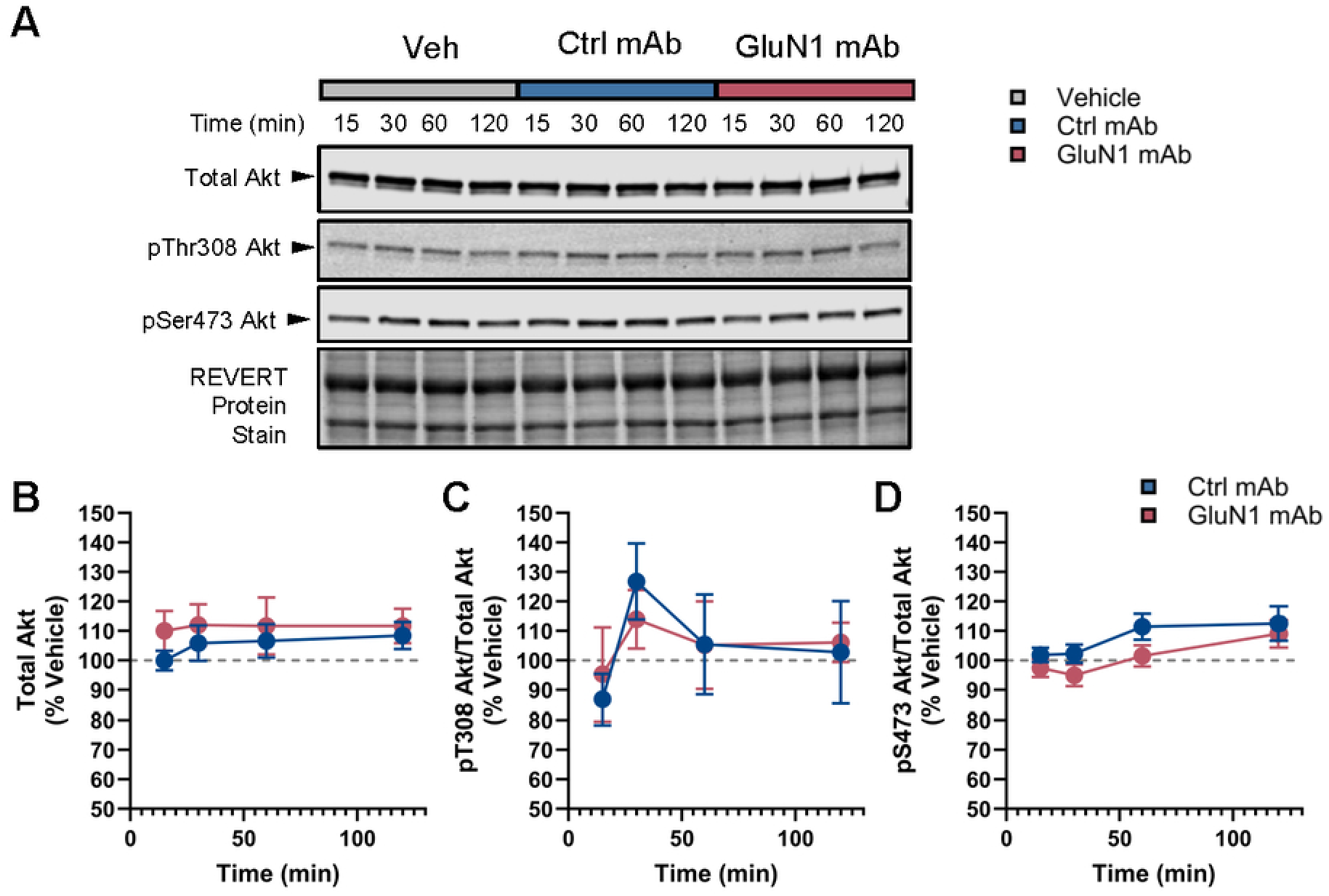
Evaluation of Akt in primary cortical neurons following human GluN1 mAb exposure. A, Representative Western blots for Akt, pThr308 Akt, and pSer473 Akt are shown from primary cortical neurons following vehicle (grey), Control mAb (blue), and GluN1 mAb (red) exposure. Timepoints are depicted. Representative image of REVERT total protein stain is also shown. B, Quantification of total Akt in the cytoplasmic fraction of primary cortical neurons following Vehicle, Control mAb, and GluN1 mAb exposure. No interaction between time and antibody specificity was detected. Antibody exposure has a statistically significant effect on total Akt level (F(2,60) = 5.28, p = 0.0078). A significant effect of human mAb on total Akt levels was detected (*p* = 0.0327). Post-hoc analyses failed to show significant effects between Ctrl mAb and GluN1 mAb conditions. n = 6 replicates. Data are normalized to vehicle at each timepoint. Plots represent mean ± SEM. C, Representative Western blots of total Akt, and two phospho-targets, pThr308 and pSer473 in the cytoplasmic fraction of primary cortical neurons after exposure to vehicle, Control mAb, and GluN1 mAb across four acute timepoints (min). C, Quantification of pT308 Akt reveals no effect of antibody specificity on pThr308 abundance. Time has a statistically significant effect on pThr308 Akt level (F(2,60) = 2.78, p = 0.0489). D, Quantification of pSer473 Akt levels across conditions reveals no effect of antibody specificity on pSer473 abundance. Time (F(2,60) = 33.88, p < 0.0001) and antibody exposure (F(2,60) = 3.88, p = 0.0260) have a statistically significant effect on pSer473 Akt levels. Post-hoc analyses failed to show significant effects between conditions. n = 6 replicates. Data are normalized to vehicle at each timepoint. Plots represent mean ± SEM. Veh, vehicle; Ctrl mAb, Control mAb.

### Kinase motif overrepresentation analysis identifies dysregulated kinase networks

We sought to explore the full complement of potential protein kinases that were altered following GluN1 mAb exposure. We utilized a comprehensive Ser/Thr kinome differential based expression analysis tool that uses differential expression of phosphopeptides to identify “hyperactivated” and “suppressed” kinases (Johnson et al. 2023). We used this tool to analyze differentially expressed phosphopeptides in the GluN1 mAb 5F5 condition compared to the Control mAb 6A condition at 2 hr and 72 hr (Fig. 6, Data S4). This analysis revealed the activities of 311 Ser/Thr protein kinases, of which 19 were significantly regulated at one timepoint. We only identified significantly hyperactivated kinases at the 2 hr timepoint (Fig. 6A). Most significantly hyperactivated kinases by GluN1 mAb 5F5 at 2 hr belong to the tyrosine-kinase like kinases (TKL) and cAMP-dependent protein kinase (PKA), cGMP-dependent protein kinase (PKG), protein kinase C (PKC) families (AGC) group. BMPR1B and BMPR1A and ALK7 are members of the TKL kinase family. These are transmembrane receptors that mediate signaling for TGF-β superfamily and bone morphogenic protein (BMP). These receptors modulate synaptic growth, maintenance, and plasticity (Kawabata, Imamura, and Miyazono 1998; Jörnvall et al. 2001; Gámez, Rodriguez-Carballo, and Ventura 2013). These activities map well onto significantly regulated GO terms following GluN1 mAb 5f5 exposure at 2 hr (Fig. 3A). GRK2, GRK3, and GRK5 are members of the AGC kinase family and phosphorylate activated G-protein coupled receptors (GPCRs) leading to desensitization. GRK2 and GRK3 belong to GRK2 subfamily, while GRK5 belongs to GRK4/5/6 subfamily. GRKs are essential for promoting recruitment of arrestins to activated GPCRs and subsequent internalization. GRK2 serves as a signaling scaffold for ARF and ERK/MAPK pathways (Pitcher et al. 1999). Together, these kinases may be hyperactivated through regulatory feedback loops and shared downstream pathways that control neuronal health, survival, and excitability following GluN1 mAb 5F5 exposure.

**Figure 6:**
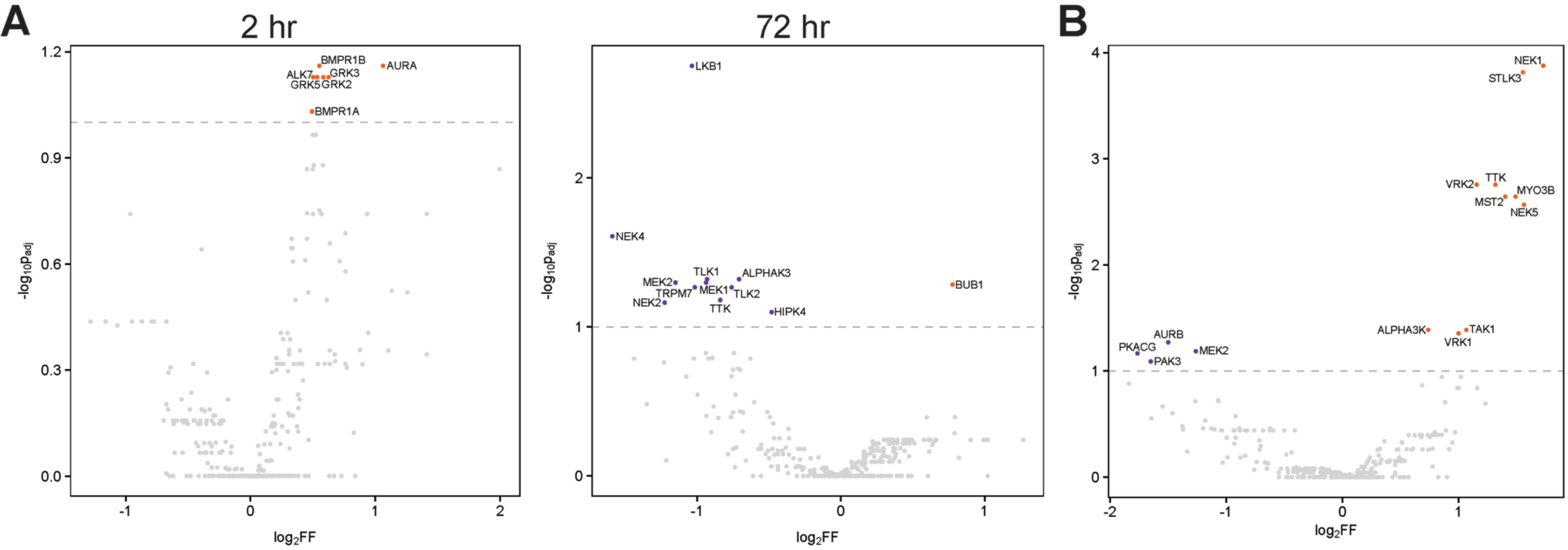
Identification of kinases whose targets are overrepresented in the GluN1 mAb- and JB2-induced phosphoproteomes. A, Volcano plots of activated or suppressed kinases in GluN1 mAb condition at 2 hr and 72 hr timepoints. Only Ser/Thr kinases were included (n = 311). B, Volcano plot of activated or suppressed kinases in NMDAR activator JB2 condition (Burgdorf et al. 2023). Activated (orange) or suppressed (purple) kinases at 1 hr timepoint are highlighted; background unregulated kinases in grey. Most kinases that are regulated by GluN1 mAb are not significantly regulated by JB2. Kinases with *p* < 0.05 and fold change > 2 were considered either activated (orange) or suppressed (purple).

As a complementary analysis, we also tested the JB2-regulated dataset (Burgdorf et al. 2023); this identified a distinct set of protein kinases. Despite the significant overlap in regulated proteins between GluN1 mAb 5F5 and NMDAR activator JB2 (Fig. 2F), there were no overlapping regulated kinase activities identified. JB2 led to hyperactivation of 10 kinases and suppression of 4 kinases (Fig. 6B). This lack of overlapping kinase activity between GluN1 mAb 5F5 and JB2 may suggest regulation of distinct signaling pathways that converge on common phosphoprotein substrates. Together, these data support the interpretation that GluN1 mAb exposure causes unique and dynamic changes to protein kinase signaling pathways downstream of NMDAR function.

We also evaluated the regulation of kinase activity by GluN1 mAb 5F5 compared to Control mAb after 72 hr exposure. At 72 hr, GluN1 mAb incubation led to overall downregulation of kinase activity, with 11 of 12 kinases suppressed compared to Control mAb 6A (Fig. 6A). Interestingly, kinases that do not fall within the traditional kinase families, referred to as “other,” comprise half of the significantly regulated kinases at 72 hr by GluN1 mAb 5F5 incubation (Fig. 6A). The kinases within this group, many of which are primarily Ser/Thr kinases, do not generally share significant similar features (Manning et al. 2002; Zhang et al. 2021). The STE family was the next most regulated group, with MEK1 and MEK2 suppressed by GluN1 mAb at 72 hr. MEK1 and MEK2 are critical components of the MAPK/ERK pathway, which is one of the most well-documented signaling pathways downstream of NMDARs. Reduced MAPK/ERK pathway signaling would be expected following NMDAR internalization. The only hyperactivated kinase activity was BUB1, a critical mitotic checkpoint kinase typically present in proliferating cells. The lack of overlap of regulated kinases between the 2 hr and 72 hr timepoints suggest dynamic regulation of kinase pathways over time following GluN1 mAb 5F5 exposure.

## Discussion

A better understanding of the molecular pathophysiology of AIE may yield novel therapeutic targets to modulate in the treatment of this grave illness. A promising approach to unravel this pathophysiology is to investigate the complex effects of the pathogenic antibodies that drive AIE. The current study investigates the specific actions of GluN1 mAb 5F5, derived from a patient with anti-NMDAR encephalitis, on synaptic protein levels and downstream molecular changes in kinase signaling networks in primary cortical neurons. Our results provide insights into the synaptic disruptions caused by GluN1 mAb and offer a window into potential novel therapeutic targets for anti-NMDAR encephalitis. These findings suggest widespread remodeling of the phosphoproteome, likely mediated through activation of the NMDAR.

Upon application of GluN1 mAb 5F5 to primary cortical neurons for 2 hr, NMDAR subunit and PSD-95 levels are unchanged compared to Control mAb 6A. Our confidence in these negative results is bolstered by our ability to detect robust changes in synaptic protein abundance in both SN and PSD compartments following NMDA stimulation. Perhaps part of the reason that Control mAb 6A regulated some phosphoproteins at 2 hr is related to the observed effect on GluN2B levels in SN fractions. Nonetheless, by focusing on synaptically enriched proteins, the current study aligns with recent research showing that NMDAR impairment can alter PSD-95 localization (Dharmasri et al. 2024) and subsequently influence synaptic strength and plasticity, which are foundational to learning and memory processes. The current study highlights that despite exposure to GluN1 mAb 5F5, NMDAR subunits and PSD-95 are unaffected, indicating a resilience of certain components within the NMDAR complex. These findings emphasize that the molecular synaptic function can be selectively altered by GluN1 mAb 5F5 without a complete dismantling of NMDAR complexes at early timepoints.

Phosphoproteomic profiling sheds light on the broader impact of GluN1 mAb 5F5 on protein phosphorylation within synaptic compartments. Similar approaches have been employed to interrogate changes in models of infectious encephalitis (Zhang et al. 2015; Ye et al. 2016; Sui et al. 2024). Although our treatment condition is highly specific, the NMDAR subunit that is targeted has widespread consequences on cellular function. Therefore, this phosphoproteomics approach has enabled us to capture both specific details of the response as well as general mechanisms of GluN1 activity. Using label-free quantitative mass spectrometry, we identified distinct phosphorylation patterns, with phosphopeptides showing altered abundance after treatment with the GluN1 mAb 5F5. The phosphoproteomic results demonstrate widespread changes in signaling pathways, particularly in proteins involved in synaptic function, neurotransmitter signaling, and microtubule dynamics. Triangulating the phosphoproteome signature we identified against archival data to infer shared regulatory processes yielded key insights about the function of the GluN1 mAb. These bioinformatic analyses reveal that the GluN1 mAb-induced phosphoproteome is positively associated with NMDAR activation and negatively associated with its inhibition. Consistent with activation of the NMDAR, one upstream regulator identified by our analysis was GSK3β. Akt is a negative regulator of GSK3β; consistent with a GSK3β activated state, we observed a non-significant trend towards decreased Akt phosphorylation by GluN1 mAb 5F5 (Fig. 5D).

Overall, our results suggest that GluN1 mAb 5F5 mediates its effect on the phosphoproteome, and perhaps on cellular function in general, by activating the NMDAR. Previous work has utilized kinase microarray of lysates from primary hippocampal neurons (Hunter et al. 2024) or proteomic analyses of hippocampal samples from passive transfer animal model (Ceanga et al. 2023) to explore molecular consequences of GluN1 antibodies. We used label-free phosphoproteomics and identified kinase activities, including CaMKII and GSK3β, as differentially regulated in SN from rat primary neurons following GluN1 mAb 5F5 exposure (Fig. 2). GSK3 was identified as potentially increased by GluN1 antibodies in a prior study based on three serine/threonine specific phosphosites (Hunter et al. 2024); however, CaMKII was not identified by the microarray kinase screen. These differences are likely to be due to our study’s use of 1) SN to target synapses versus lysates from whole neurons, and 2) phosphoproteomics that is more sensitive than microarray kinase assay.

Our analyses also reveal GluN1-dependent remodeling of numerous synapse-related pathways, particularly those related to neurotransmitter trafficking; cytoskeleton regulation; and signal transduction, such as that associated with calmodulin binding. Calmodulin regulates numerous synaptic processes, including CaMKII and other signaling proteins. Its perturbation may explain some of the observed phosphorylation changes, especially since calmodulin is closely tied to calcium signaling dynamics within neurons. CaMKII is pivotal for synaptic potentiation, and its dysregulation can disrupt memory encoding and retrieval processes (Lisman, Yasuda, and Raghavachari 2012). Interestingly, although pharmacologic NMDAR activation translocated CaMKII to the PSD, GluN1 mAb 5F5 did not alter basal or stimulated levels of CaMKII, suggesting that CaMKII recruitment to the PSD is preserved in the presence of GluN1 mAb 5F5 at early timepoints. The alterations in CaMKII phosphorylation map to the variable linker region of the molecule (Fig. 4B), suggesting CaMKII holoenzyme formation or protein-protein interactions may be affected. Indeed, phosphorylation of Thr337 CaMKIIα has been implicated previously in regulating the association of CaMKII with GluN2B (Wang et al. 2017). Indeed, structural and functional properties of CaMKII have been shown to be critical in NMDAR regulation of synaptic plasticity (Incontro et al. 2018). Further studies are needed to explore the detailed mechanisms at play in the regulation of CaMKII structure and function following GluN1 mAbs.

These findings offer a potential mechanism by which GluN1 antibodies might contribute to the clinical symptoms observed in anti-NMDAR encephalitis. By disrupting key signaling pathways, these antibodies alter intracellular kinase signaling that is responsible for the cognitive deficits, seizures, and neuropsychiatric symptoms characteristic of the disorder. The observed dysregulation of calmodulin and ARF protein signaling may serve as biomarkers or therapeutic targets for treating the disease. Our findings of altered synaptic function without NMDAR loss align with recent studies that demonstrate direct channel regulation by GluN1 antibodies (Michalski et al. 2024) and expanded epitopes of GluN1 antibodies (Michalski et al. 2024; Wang et al. 2024). Rather than promoting surface NMDAR levels, future treatments could focus on modulating disrupted signaling cascades to stabilize synaptic structure and function. Such work would expand the concept that GluN1 antibody-mediated kinase signaling is a core molecular pathophysiological feature of the disease. Future studies should investigate if interventions aimed at stabilizing kinase networks can mitigate the effects of GluN1 antibodies.

The use of phosphoproteomics provides unique insights into the GluN1 antibody-induced changes within synaptic compartments. However, there are several limitations in this approach. For example, the reliance on this *in vitro* model system may not fully capture the *in vivo* complexity of synaptic signaling in anti-NMDAR encephalitis. Most of the phosphopeptides identified in our dataset were phosphoserine residues, consistent with the most common phosphorylation site in the mammalian proteome being Ser (Kalyuzhnyy et al. 2022). Nonetheless, this biased the evaluation of downstream kinases and pathways to primarily phosphoserine, and more specifically to proline-directed protein kinases, in our dataset. Therefore, our ability to fully interrogate protein phosphorylation changes, particularly at phosphothreonine or phosphotyrosine signaling pathways, is limited. We elected to directly compare between GluN1 mAb 5F5 and Control mAb 6A datasets. The rationale for this decision was to avoid misattributing nonspecific effects of the vehicle conditions to human mAb. Indeed, we observed many effects of the Control mAb 6A compared to vehicle condition. Possible explanations could include 1) off-target low-affinity antibody binding, 2) interaction with targets in the growth media, or 3) through Fc portions of the antibody. Indeed, recent evidence supports that Fc glycosylation status may have diverse impact on antibody effects in neurons (Terroba-Navajas et al. 2024). We also observed an effect of time on our dataset as revealed by PCA (Figure S2D), which was particularly evident in vehicle conditions at the two timepoints. Hence, our decision to focus on pathways that were regulated uniquely by GluN1 5F5 compared to Control mAb was directed at preventing exclusion of disease-relevant pathways. Addressing these limitations in future research will improve our understanding of how these findings translate into anti-NMDAR disease mechanisms.

In summary, we elucidate the impact of GluN1 antibodies on cortical neuron signaling by examining protein phosphorylation and kinase pathway dysregulation. Through a targeted phosphoproteomic approach, it highlights specific receptor subunits, synaptic proteins, and pathways, such as calmodulin and ARF protein signaling, affected by GluN1 antibodies. Critically, our study provides proteome-level evidence that the pathological GluN1 mAb 5F5 functions by activating the NMDAR. Here, the study advances our knowledge of the molecular neurobiology underlying AIE in relation to synaptic stability, pathway disruption, and potential therapeutic targets. By expanding our understanding of signaling regulators, the study provides a foundation for potential therapeutic strategies that could stabilize or restore disrupted signaling pathways in anti-NMDAR encephalitis. These findings deepen our understanding of how GluN1 antibodies affect neural structure and function and highlight the importance of focusing on pathway modulation as a treatment strategy for AIE.

## Conflict of interest statement

The authors declare no competing financial interests.

## Acknowledgements

This work was supported by NIH K08NS114039-01 and seed funds from University of Maryland School of Medicine (D.R.B.). The authors thank members of the Blanpied laboratory for critical discussions in the preparation of this manuscript. We acknowledge the support of the University of Maryland, Baltimore, Institute for Clinical & Translational Research (ICTR) and the National Center for Advancing Translational Sciences (NCATS) Clinical Translational Science Award (CTSA) and the Program for Research Initiated by Students and Mentors (PRISM), UMSOM Office of Student Research.

Current affiliations are School of Chemistry and Molecular Biosciences, The University of Queensland, St. Lucia, Australia (W.H.); and Solomon H. Snyder Department of Neuroscience, Johns Hopkins University School of Medicine, Baltimore, MD, USA (N.G.).

## Supplemental Figures

**Figure S1:**
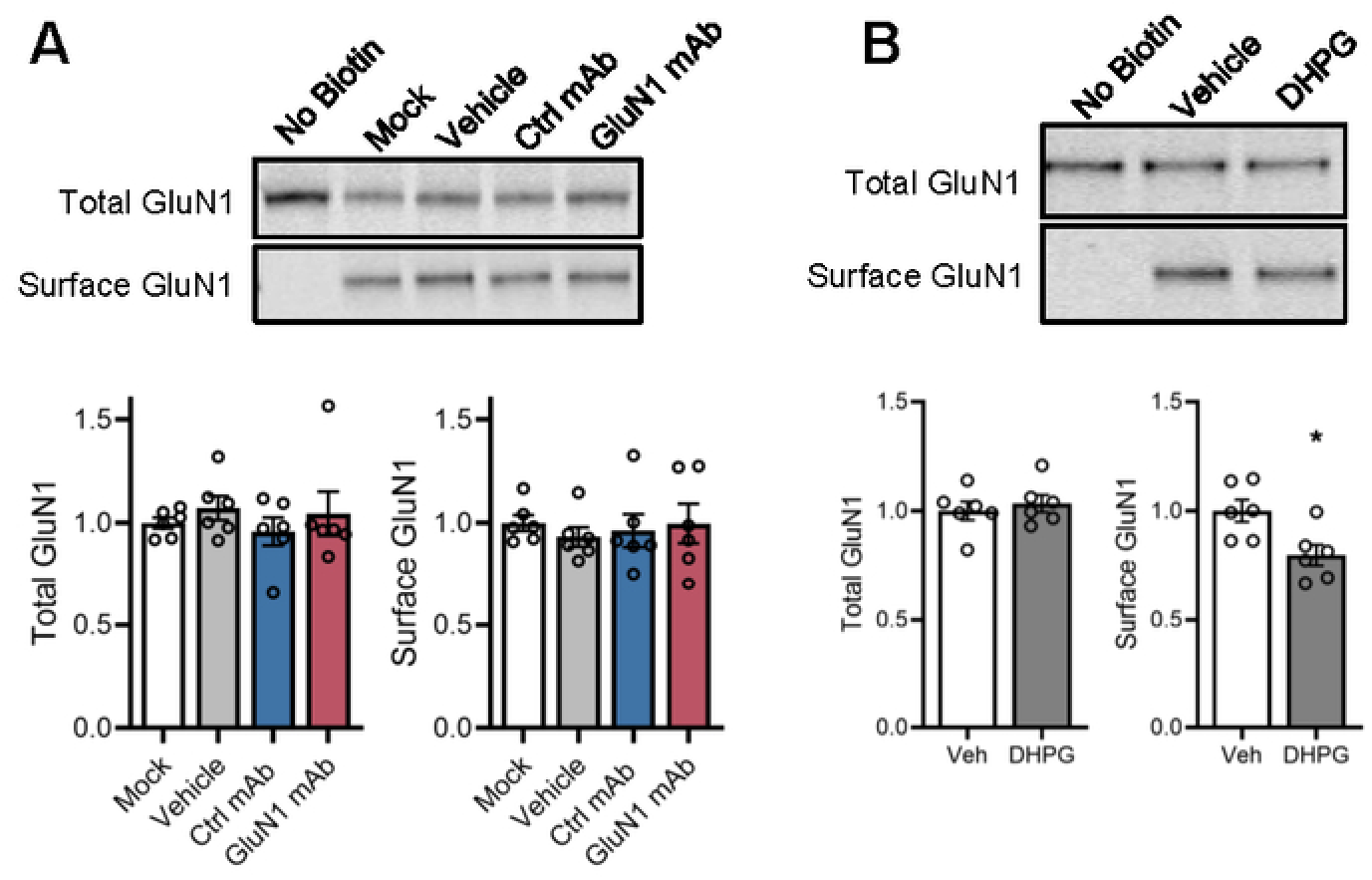
No effect of mAb exposure on surface GluN1 levels by surface biotinylation assay. A, Representative blots for GluN1 are shown. No biotin condition demonstrates the specificity of assay for surface targets. Histogram depicts normalized surface and total GluN1 levels. mAb had no effect on surface GluN1 levels (F(3,20 = 0.2108, *p* = 0.8877, one-way ANOVA)). n = 6 per group. Data are mean ± SEM. B, DHPG (100 μM) for 5 min, followed by 60 min washout prior to surface biotinylation caused reduction of surface GluN1 levels in primary cortical neurons compared to vehicle (*p* = 0.0157). n = 6 per group. Representative blots for GluN1 are shown. n = 6 per group. Data are mean ± SEM, with individual replicates shown. * *p* < 0.05, unpaired Student’s t-test. SN, synaptoneurosome; PSD, postsynaptic density; Veh, vehicle; DHPG, (*RS*)-3,5-dihydroxy phenylglycine.

**Figure S2:**
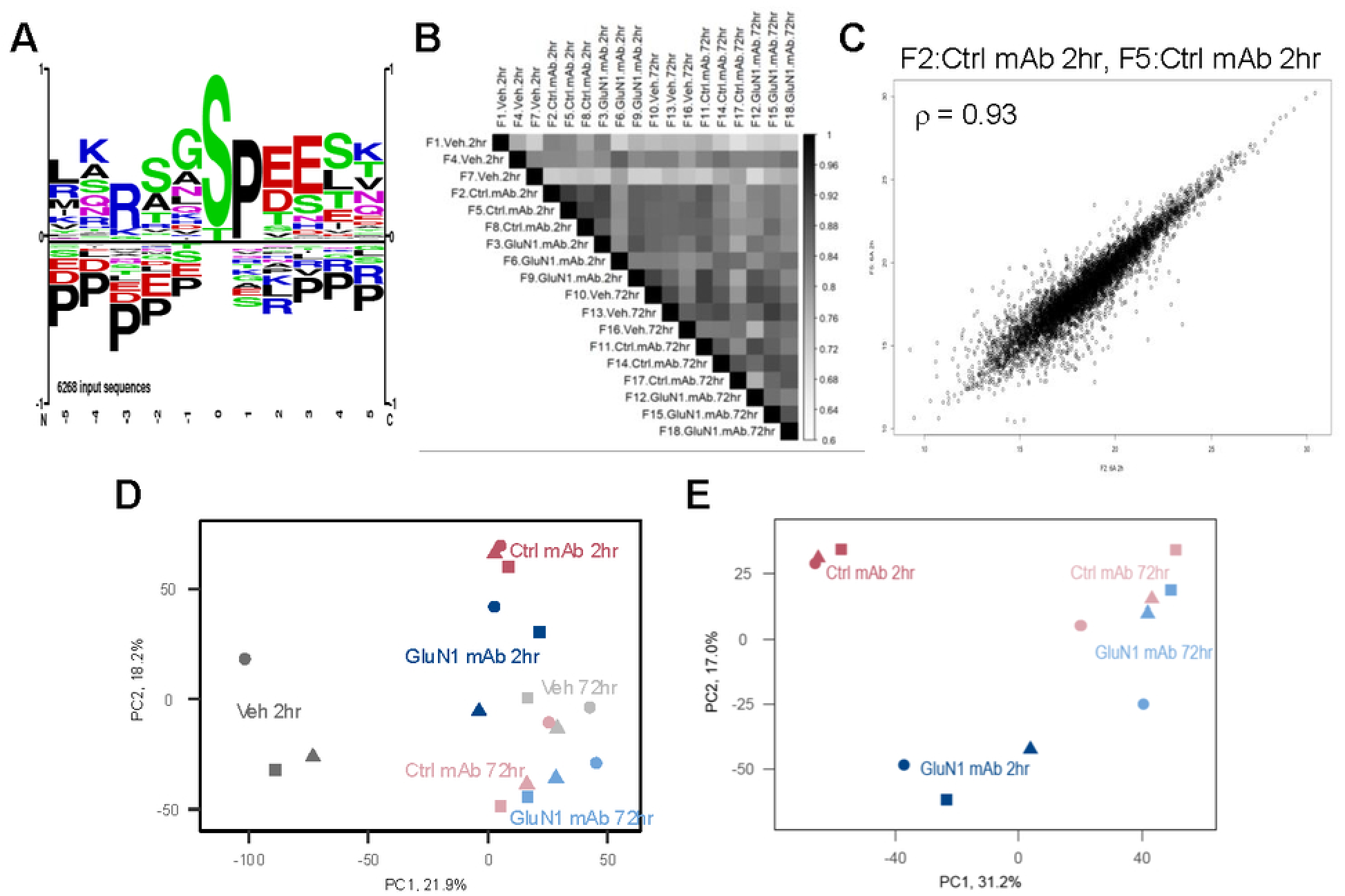
Evaluation of phosphoproteomic dataset across conditions and time points. A, Motif analysis of all phosphopeptides with identified phosphorylation sites demonstrates that the most highly overrepresented motifs correspond to proline-directed Ser and Thr residues. B, Spearman’s rank correlations across all replicates, conditions, and timepoints. A Spearman’s rank correlation indicated that there was a significant and strong positive relationship between groups. C, The highest Spearman’s ‘Rho’ correlation was between F5: Ctrl mAb 2 hr and F8: Ctrl mAb 2 hr, *ρ* = 0.93, *p* = 2.2e^-16^. D, Principal component analysis plot comparing all conditions. PC1 and PC2 are shown on the x and y axis, comprising a total of 40.1% of the variance across the conditions and replicates. Conditions are represented by color, while replicates are represented by shape. There are a total of n = 4080 phosphopeptides included across treatment groups (treatment = Veh, Ctrl mAb, GluN1 mAb, timepoint = 2 hr, 72 hr), with n = 3 replicates. E, Principal component analysis plot comparing GluN1 mAb and Control mAb conditions over time. PC1 and PC2 are shown on the x and y axis, comprising a total of 48.2% of the variance across the conditions and replicates.

Data S1: Overview of Phosphoproteomic Data for GluN1 mAb vs Control mAb vs Vehicle

Data S2: Overview of Top Upstream Regulators for GluN1 mAb vs Control mAb

Data S3: Gene Ontology (GO) Enrichment Analysis for GluN1 mAb vs Control mAb

Data S4: Kinase Motif Enrichment Analyses

## Notes

### Competing Interest Statement

The authors have declared no competing interest.

